# A load-bearing function for the cytoplasmic membrane of *Escherichia coli*

**DOI:** 10.1101/2025.10.02.680147

**Authors:** Jiawei Sun, Miriam Min-Chen Cheng, Ellen van Wijngaarden, Christopher J. Hernandez, Thomas G. Bernhardt, Kerwyn Casey Huang

**Affiliations:** Department of Bioengineering, Stanford University, Stanford, CA 94305, USA; Sibley School of Mechanical and Aerospace Engineering, Cornell University, Ithaca, NY 14850, USA; Departments of Orthopaedic Surgery and Bioengineering and Therapeutic Sciences, UCSF; Department of Microbiology, Harvard University Medical School, Boston, MA 02115, USA; Howard Hughes Medical Institute, Harvard University Medical School, Boston, MA 02115, USA; Department of Microbiology and Immunology, Stanford University School of Medicine, Stanford, CA 94305, USA; Chan Zuckerberg Biohub, San Francisco, CA 94158, USA

**Keywords:** MreB, cytoplasmic membrane, cell wall, peptidoglycan, mechanical coupling, osmotic shocks, membrane stiffness, elongasome, transglycosylase, MltG

## Abstract

The structural integrity of bacterial cells is traditionally attributed to the peptidoglycan cell wall, and more recently to the outer membrane, with the cytoplasmic membrane assumed to be mechanically passive. Cells lacking filaments of the actin homolog MreB are more bendable, suggesting a role for the cytoskeleton in cell stiffness. Here, we show that MreB does not stiffen the envelope directly, but instead mechanically couples the cell wall to the cytoplasmic membrane through its role in peptidoglycan synthesis, increasing resistance to bending. Under hyperosmotic stress, MreB relocalized to the poles, forming linkages that prevent membrane detachment from the cell wall and attenuate cytoplasmic contraction. Disruption of MreB filament formation, nutrient starvation, or inactivation of glycan elongation factors abolished or reduced this coupling, revealing that peptidoglycan biosynthesis actively mediates stress distribution across surface layers. Our findings redefine the bacterial envelope as a mechanically integrated composite, with the cytoplasmic membrane having substantial load-bearing capacity.

## Introduction

The bacterial cell envelope is a versatile and dynamic structure that acts as a chemical barrier and supports cell growth and division. The envelope bears mechanical stresses from both inside the cell and the external environment^1^, including forces due to fluid flow, solid substrates, and osmolarity changes^2^. In particular, a major function of the envelope is to withstand the turgor pressure generated by high solute concentrations in the cytoplasm^3,4^. In Gram-negative bacteria, the envelope consists of a peptidoglycan cell wall sandwiched by the cytoplasmic and outer membranes. The outer membrane is an asymmetric bilayer with phospholipids in the inner leaflet and lipopolysaccharides in the outer leaflet that have important roles in chemical permeability^5^. The cell wall is a rigid macromolecule that has well-recognized importance as an antibiotic target^6^ and in resisting cell expansion due to turgor pressure^7^. Robust envelope growth and division requires coordination by a host of enzymes^8^. In many rod-shaped bacteria, an essential component of the elongation machinery is the actin homolog MreB, which forms filaments that bind to the inner surface of the cytoplasmic membrane and interact with the cell-wall synthesis machinery to control cell shape through the insertion pattern of peptidoglycan precursors^9^.

The cytoplasmic membrane is often ignored in discussions of bacterial mechanics. Nonetheless, previous studies hinted at an underappreciated load-bearing capacity of the cytoplasmic membrane in resisting expansion due to turgor pressure. The gain-of-function mutation *mlaA\** in *E. coli* results in rapid transfer of phospholipids from the cytoplasmic membrane to the outer membrane during starvation, and this transfer causes the cytoplasmic membrane to shrink by ∼20% and detach from the cell wall^10,11^. Once the cytoplasmic membrane of starved *mlaA\** cells ruptures, its contents expand in volume to reach approximately the original cell length pre-plasmolysis^10,11^. The ability of these cells to persist in a shrunken state for several minutes, together with the subsequent expansion, suggests that the cytoplasmic membrane alone can bear turgor pressure upon material loss and hence has substantial load-bearing capacity.

Recent studies have revealed that many other components of the envelope beyond the cell wall can impact cellular mechanical integrity^12^. The outer membrane has stiffness properties similar to those of the cell wall and is critical for maintaining cell viability during rapid changes in external osmolarity^13^. Mechanical linkages between the cell wall and outer membrane such as Braun’s lipoprotein (Lpp) and OmpA strengthen the cell envelope and determine periplasmic width^14–17^. Mechanosensitive channels in the cytoplasmic membrane enable rapid osmolyte efflux to protect cells against hypoosmotic shocks^17,18^. Beyond the envelope itself, deletion of a number of other proteins can impact the stiffness of *E. coli* cells^19^. In particular, disruption of MreB filaments by the antibiotic A22 decreased the resistance of *E. coli* cells to bending^20^. One interpretation of this finding is that MreB forms an ultrastructure that directly increases the mechanical stiffness of the envelope, mimicking the function of its eukaryotic counterpart actin^21^. However, the mechanism by which MreB confers increased bending rigidity could involve its interaction partners within the cell-wall synthesis machinery including MreC, MreD^22^, RodA, and RodZ^23,24^. During wall synthesis, MreB filaments translate approximately along the circumferential direction^25–28^. Given this motion, it is currently unclear how mechanical forces could be transmitted from MreB, bound to the inner surface of the cytoplasmic membrane, to the rest of the cell envelope.

In this study, we investigate the mechanical contribution of MreB to the cell envelope by quantifying cell wall deformation during hyperosmotic shocks. Under moderate shocks that are sufficient to cause plasmolysis, depolymerization of MreB by A22 did not significantly increase cell wall deformation, suggesting that MreB does not directly contribute to envelope stiffness. However, under large-magnitude oscillatory hyperosmotic shocks, cytoplasmic contraction gradually decreased in cells with intact MreB, whereas A22-treated cells with depolymerized MreB maintained the same degree of plasmolysis in each successive shock, indicating that MreB filaments are required to prevent cytoplasmic contraction from the cell wall. Fluorescence microscopy revealed that MreB relocalizes to the cell poles during hyperosmotic conditions, where it drives the formation of structural linkages between the cytoplasmic membrane and the cell wall that enable polar wall synthesis. Perturbations of certain cell-wall synthesis factors altered the coupling, indicating that the elongation machinery forms a physical linkage between the cytoplasmic membrane and the peptidoglycan. Using a mutant with a weak outer membrane, we show that wall-less spheroplasts can transiently remain viable even after the outer membrane ruptures, indicating that the cytoplasmic membrane can bear substantial mechanical stress. Based on these results, we propose that rather than directly strengthening the cell wall, MreB-directed cell-wall synthesis mechanically couples the cytoplasmic membrane and the cell wall, enabling the intrinsic stiffness of the cytoplasmic membrane to contribute to the overall strength of the envelope.

## Results

### Disruption of MreB polymerization does not alter cell-wall stiffness

The previous observation that A22 treatment decreases the bending rigidity of *E. coli* cells^20^ on a short time scale before transcriptional reprogramming could occur suggested that MreB might directly impact envelope stiffness. To test this hypothesis, we queried whether MreB polymerization and membrane binding affects other morphological responses to perturbations beyond cell bending. First, we reasoned that if MreB filaments were stress bearing, then their depolymerization would alter the force balance across the cell envelope and hence cell size would change. We quantified the dimensions of cells during steady-state growth in a microfluidic flow cell (Methods) with high temporal resolution before and after addition of 100 µg/mL A22, which rapidly disrupted MreB polymerization as expected (Movie S1). Cell width and length were unaffected (Fig. 1a), indicating that MreB was not bearing levels of longitudinal or circumferential stress relevant to cell morphology.

**Figure 1:**
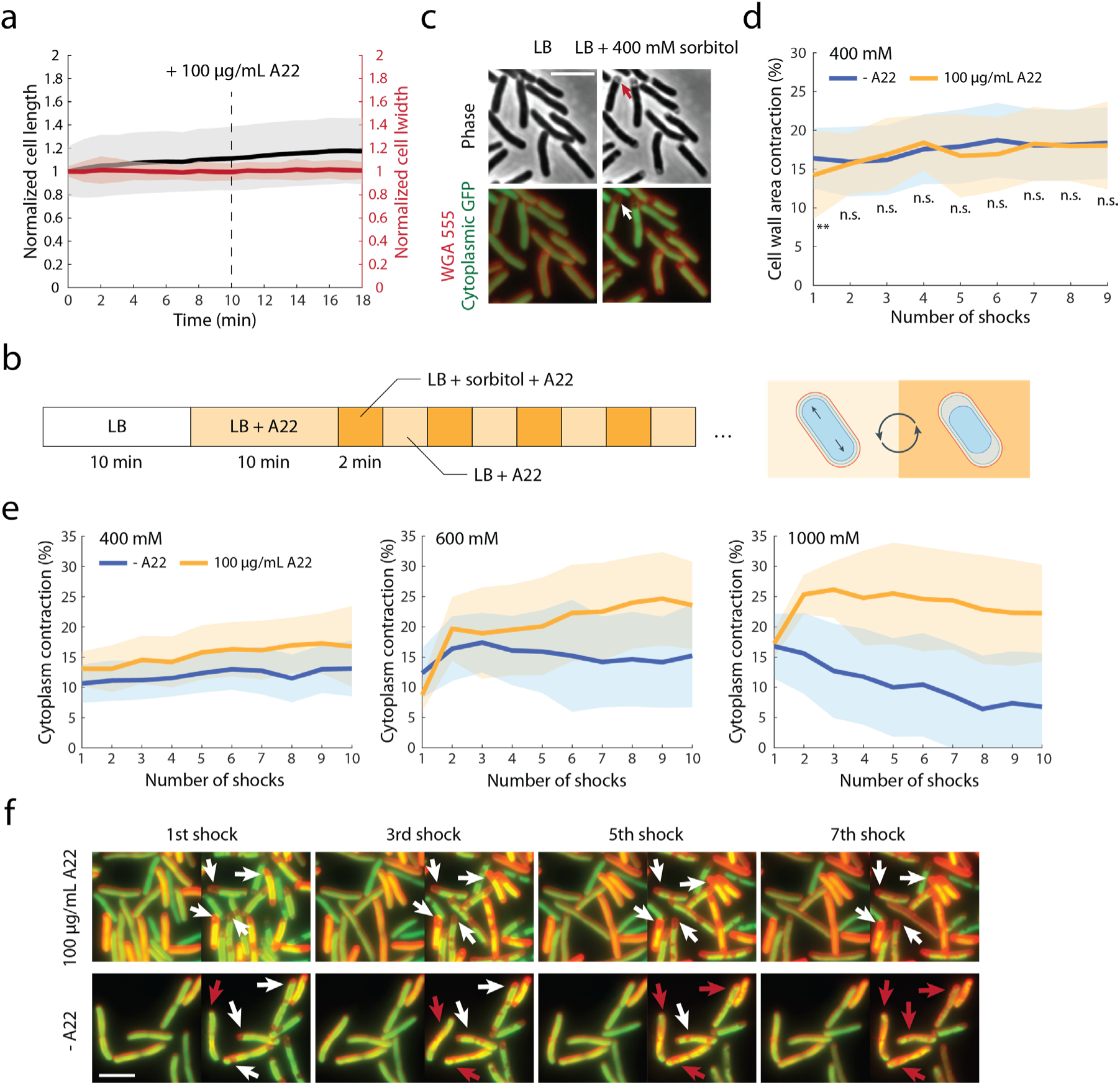
MreB couples the cytoplasmic membrane to the cell wall. a) Disruption of MreB filaments with 100 µg/mL A22 in LB medium did not alter cell length or width, suggesting that MreB filaments are not under stress during exponential growth. A22 treatment started at *t* = 10 min. b) Schematic of the oscillatory hyperosmotic shock assay in which cells were periodically subjected to hyper- and normal osmolarity (Methods). Under normal osmolarity, the cell wall is under tension due to turgor pressure; under hyperosmolarity, the cytoplasm loses water and detaches from the cell wall. c) Exposure to 400 mM hyperosmotic shock cycles led to cytoplasmic contraction, typically seen at the cell poles (red/white arrows highlight one example). The OM and the cytoplasm are labeled with wheat germ agglutinin (WGA) AlexFluor-555 and GFP expressed from a plasmid, respectively. Scale bar: 5 µm. d) Disruption of MreB filaments via pre-treatment with 100 µg/mL A22 did not alter cell wall contraction (Methods) under 400 mM oscillatory hyperosmotic shocks. **: *p* < 0.01, n.s.: *p* > 0.01; Welch’s t-test was performed for each osmotic shock cycle. e) Inhibition of cytoplasmic contraction by MreB in plasmolyzed cells was dependent on the number and magnitude of the oscillatory osmotic shocks. The impact of A22 pre-treatment on cytoplasmic contraction was prominent only under large hyperosmotic shocks (600 mM and 1 M). f) The presence of MreB filaments resulted in mechanical coupling between the cytoplasmic membrane and the cell wall (red arrows); disruption of MreB by A22 allowed unhampered plasmolysis (white arrows). Cells are labeled with WGA AF-555 and cytoplasmic GFP expressed from a plasmid. Scale bar: 5 µm.

Nonetheless, it remained possible that MreB contributes to envelope stiffness by bearing stress when cells experience mechanical perturbations that disrupt steady-state growth. To investigate this possibility, we applied oscillatory osmotic shocks in a microfluidic flow cell and measured the deformation of the cell wall upon each hyperosmotic shock in the absence and presence of A22 (Methods). In this assay, log-phase cells were alternately subjected to hyperosmotic (LB+400 mM sorbitol) and normal (LB) osmotic conditions (Fig. 1b). Each hyperosmotic shock led to contraction of the cell wall and was sufficiently large in magnitude to reduce turgor pressure to zero, as evidenced by plasmolysis of the cytoplasm (Fig. 1c). The stiffness of the cell envelope is related to the degree of deformation upon removal of turgor^12^. As observed in previous studies with 100-mM shocks^7^, the magnitude of length and width contractions was consistent across shock cycles (Fig. 1d). We found that the degree of contraction was not significantly different between cells without or with 10 min of treatment with 100 µg/mL A22 prior to the start of the shocks (Fig. 1d). Thus, disruption of MreB polymerization does not affect the mechanical response of the cell envelope to forces that induce contraction/extension.

Taken together, these results indicate that MreB does not contribute directly to cell envelope stiffness, despite the significant impact of A22 on bending rigidity^20^.

### MreB promotes a physical coupling between the cytoplasmic membrane to the cell wall

Since MreB and its associated components of the cell-wall synthesis machinery form a transmembrane structure connecting the cytoplasmic membrane to the cell wall^29^, we reasoned that MreB may prevent detachment of the cytoplasmic membrane from the cell wall during osmotic shocks. To accentuate the impact of A22 treatment, we increased the magnitude of the osmotic shock oscillations to 1 M (Methods), which substantially inhibited *E. coli* cell growth (Fig. S1). To counteract the induction of cell division by hyperosmotic shocks^30^, we also treated cells with the antibiotic cephalexin (50 µg/mL), which targets the key division protein PBP3^31^, in the microfluidic flow cell for 40 min prior to the shocks (Methods). In cells pre-treated with A22 for 10 min, the degree of cytoplasmic contraction was consistently large over the course of 10 osmotic shock cycles (Methods, Fig. 1e). In contrast, in untreated cells, the degree of cytoplasmic contraction gradually decreased over the cycles (Fig. 1e). During the last few shocks, many untreated cells exhibited little plasmolysis (Fig. 1f), suggesting that polymerized MreB gradually inhibited detachment of the cytoplasm from the cell wall. In addition to inhibiting detachment of the cytoplasm from the cell wall, the linkages mediated by MreB also caused the cell wall to buckle under the negative pressure generated by cytoplasmic contraction for 1 M shocks, resulting in thinning of the cylindrical region around midcell (Fig. S2). Treatment with 80 µg/mL MP265, a structural analog of A22, led to similar effects as A22 (Fig. S3, Methods), arguing against potential off-target effects of A22. These findings indicate that MreB promotes physical coupling of the cytoplasmic membrane to the rest of the cell envelope.

### Plasmolysis results in relocalization of MreB to the cell poles

To probe the dependence of plasmolysis dynamics on oscillatory osmotic shock magnitude, we tracked the subcellular localization of MreB in a strain expressing an MreB-msfGFP sandwich fusion that complements *mreB* deletion^32,33^ (Table S1). During steady-state growth, MreB filaments localized primarily to the cylindrical region of the cell (Fig. 2a)^33^, as expected, where they guide the insertion of new cell wall material^27^. During 1 M oscillatory osmotic shocks, the first few hyperosmotic shocks induced cytoplasmic contraction mostly near the cell poles (Fig. 2a), indicating the lack of coupling between the cytoplasmic membrane and the rest of the envelope at the poles, where MreB was largely absent. However, in subsequent osmotic shock cycles, MreB partially relocalized to the cell poles, resulting in a more homogeneous distribution of MreB across the cell surface (Fig. 2a-c). This gradual relocalization coincided with the decrease in cytoplasmic contraction (Fig. 1e). In contrast, lower-magnitude (400 mM) osmotic shocks for which cytoplasmic contraction remained constant over shock cycles (Fig. 1e) resulted in less relocalization of MreB (Fig. 2c). Thus, we reasoned that repeated, large-magnitude osmotic shocks led to the formation of MreB-mediated linkages between the cytoplasmic membrane and the other envelope layers at the cell poles that are stable over the 4-min period of the shock cycles, which limited contraction during subsequent shock cycles.

**Figure 2:**
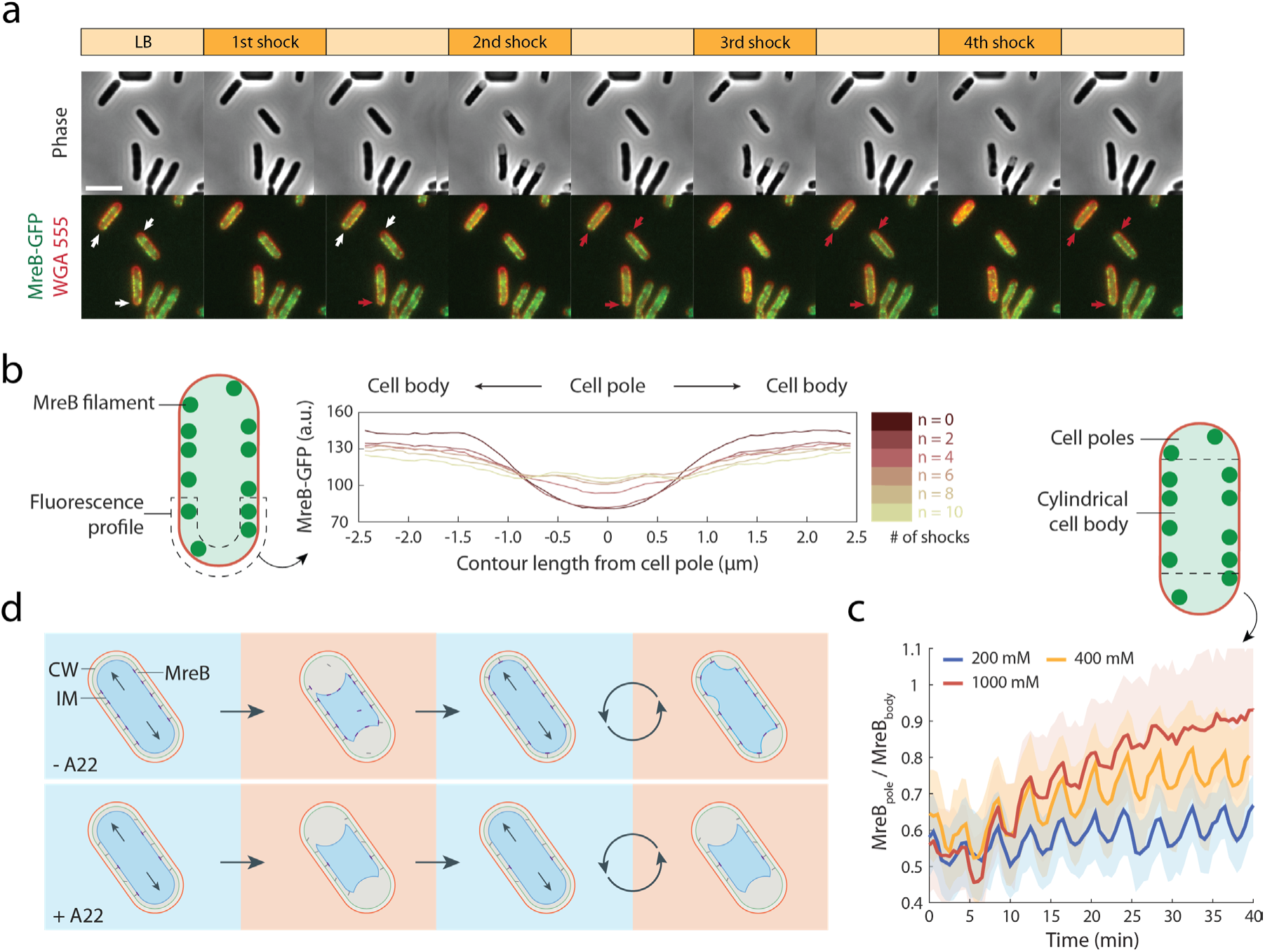
Hyperosmolarity causes relocalization of MreB filaments to the poles. a) 1-M oscillatory osmotic shocks resulted in relocalization of MreB-msfGFP filaments to cell poles (red arrows), which are generally depleted of MreB-msfGFP during normal growth (white arrows). Scale bar: 5 µm. b) MreB-msfGFP profiles of individual cell poles, averaged across the contours of *n*=348 cells, demonstrate the increasingly homogenous distribution of MreB after more 1-M hyperosmotic shocks. The profiles span the pole and surrounding region (left schematic). c) The increase in the ratio of MreB-msfGFP intensity at the cell poles compared to the cylindrical cell body (left schematic) demonstrates the relocalization of MreB to the poles. The degree of relocalization was dependent on the shock magnitude. The sawtooth behavior is due to repeated plasmolysis and subsequent recovery. d) Schematic of MreB relocalization under hyperosmotic shocks and formation of new mechanical linkages at the cell poles.

To test whether hyperosmotic shocks themselves trigger MreB relocalization, we varied the duration of hyperosmolarity and observed nearly identical relocalization trajectories over assay time regardless of the number of shock cycles (Fig. S4). These results indicate that MreB filaments relocalize continuously during plasmolysis and recovery, reflecting a process that depends on the time under osmotic perturbation rather than an acute effect of the shock.

Taken together, these findings suggest that hyperosmolarity drives relocalization of MreB to the poles, where they appear to establish new physical linkages between the cytoplasmic membrane and cell wall that limit cytoplasmic contraction during subsequent osmotic shocks (Fig. 2d).

### Peptidoglycan elongation couples the cytoplasmic membrane and cell wall

Since MreB resides solely on the inner surface of the cytoplasmic membrane, we surmised that any physical connection to the cell wall must rely on other cellular components. While many proteins interact with both the cytoplasmic membrane and cell wall, presumably only a subset are critical for maintaining the overall physical coupling between layers. We hypothesized that the most likely candidates would be components of the cell-wall synthesis machinery. Supporting this hypothesis, when we performed oscillatory osmotic shocks with 1 M sorbitol in a nutrient-free buffer osmotically balanced with LB (0.85X PBS, Methods), regardless of A22 treatment, cells exhibited plasmolysis without a reduction in cytoplasmic contraction over time (Fig. 3a). These data suggest that linkage between the membrane and cell wall requires cell growth. Repeated osmotic shocks in the nutrient-free media still resulted in MreB relocalization to the poles (Fig. S5), indicating that MreB relocalization is not dependent on wall synthesis and is not sufficient for linking the cytoplasmic membrane to the cell wall.

**Figure 3:**
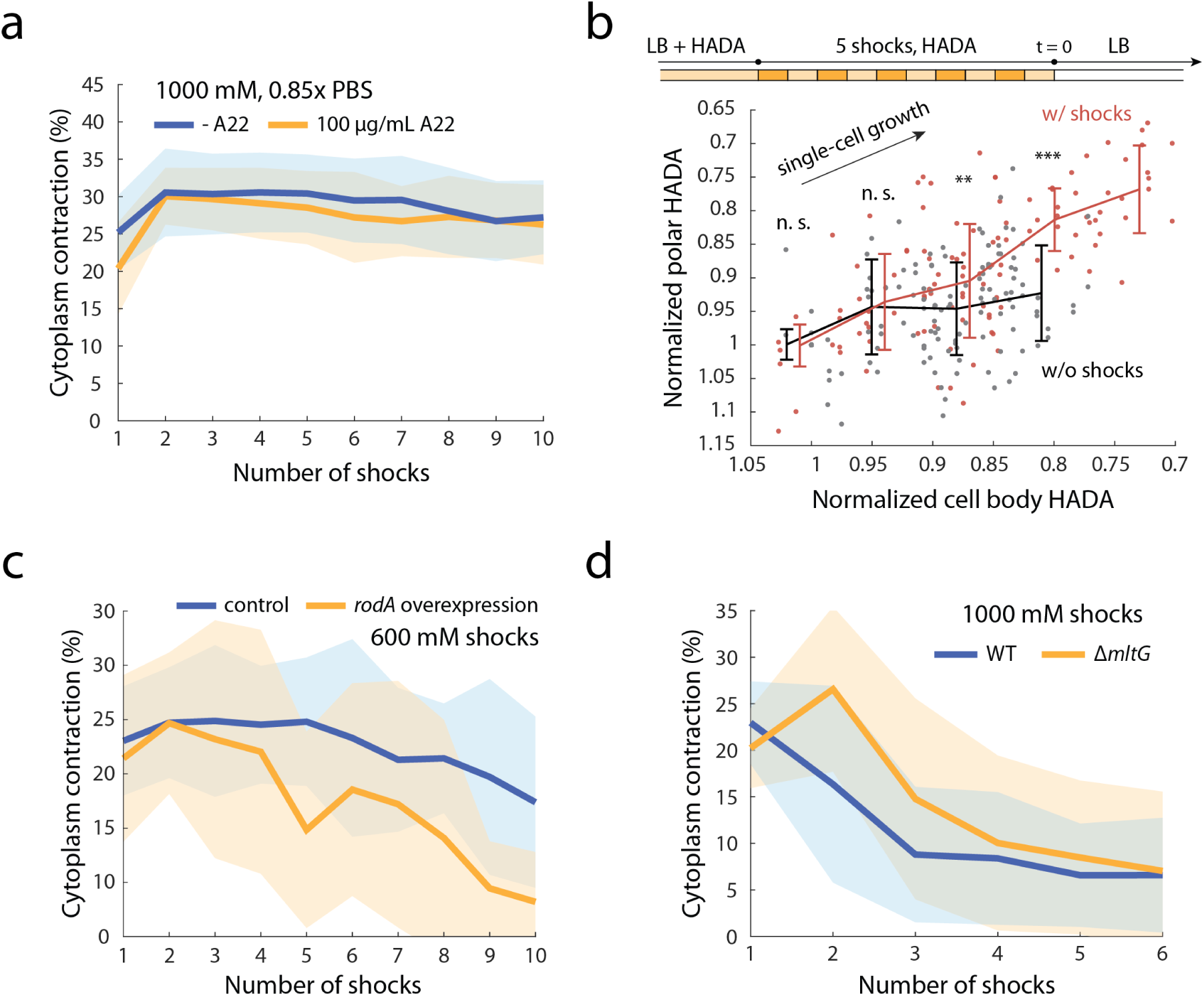
Inactivation of cell-wall synthesis alters the coupling between the cytoplasmic membrane and the peptidoglycan. a) In 0.85X PBS, wild-type cells exhibited plasmolysis without any reduction in cytoplasmic contraction over repeated 1-M osmotic shocks, regardless of A22 treatment. b) Pulse-chase experiment in cells pre-labeled with 1 mM HADA. Compared to cells grown without shocks, cells exposed to shocks exhibited greater decreases in fluorescence at the poles, when evaluated at the same level of fluorescence decay in the cell body, indicating increased polar peptidoglycan synthesis following osmotic shock-induced MreB relocalization. Top: schematic of the HADA labeling and hyperosmotic shock protocol. Cells were tracked in a microfluidic flow cell during growth without HADA starting at *t*=0. Each data point corresponds to an individual cell pole. *N*≥25 cells were tracked for each condition. **: *p* < 0.01, ***: *p* < 0.001, n.s.: *p* > 0.01; Welch’s t-test was performed on binned data points. c) Overexpression of RodA reduced cytoplasmic contraction during 600-mM osmotic shocks. d) Δ*mltG* cells exhibited a slower transition to the coupled state compared to wild type during 1-M shocks. The degree of contraction was significantly different (Welch’s t-test: *p* < 0.01) in the first 3 shocks and similar after *n*>3 shocks.

To test whether MreB relocalization to the poles leads to polar wall synthesis, rather than the side-wall synthesis typical of normal growth, we performed pulse-chase experiments in cells labeled with 3-[[(7-Hydroxy-2-oxo-2H-1-benzopyran-3-yl)carbonyl]amino]-D-alanine hydrocholoride (HADA), which is inserted into the peptidoglycan at locations of growth^33,34^. During subsequent growth in HADA-free medium (Methods), the ratio of fluorescence at the poles relative to the cell body decayed more rapidly in cells undergoing hyperosmotic shocks, indicating increased polar wall synthesis (Fig. 3b). Thus, polar localization of MreB leads to relocalization of elongation machinery function.

We next hypothesized that proteins involved in cell-wall synthesis^29^ contribute to the membrane-cell wall connection. To identify such factors, we performed oscillatory shocks in combination with genetic or chemical perturbation of elongation-associated targets. First, we overexpressed RodA from an inducible plasmid (Table S1) prior to osmotic shocks (Methods). Opposite to depolymerization of MreB by A22, cytoplasmic contraction was reduced during repeated plasmolysis compared with no induction (Fig. 3c). Overexpression of a dominant-negative mutant *rodA^D262N^* (Table S1), which blocks the glycosyltransferase activity of RodA^35^, led to a similar phenotype (Fig. S6). These results suggest that increasing the copy number of RodA is sufficient to enhance the coupling between MreB and the cell envelope, regardless of RodA activity.

To assess the contribution of other elongation-associated PBPs, we inhibited their activity using β-lactam antibiotics, which specifically target transpeptidase function. Inhibition of PBP1A/B by cefsulodin, PBP2 by mecillinam, or all PBPs by ampicillin had no effect on cytoplasmic contraction during the shocks (Fig. S7), suggesting that transpeptidase function is not required for establishing the cytoplasmic membrane–cell wall connection.

To probe the role of glycan strand synthesis, we focused on a deletion mutant of *mltG* (Table S1), which encodes a cytoplasmic membrane-anchored enzyme implicated in the termination of peptidoglycan synthesis^36^. Deletion of *mltG* is predicted to enhance the association of the elongation machinery with the cell wall and results in longer glycan strands^36^. Under oscillatory osmotic shocks, Δ*mltG* cells exhibited a slower transition to the state of lower cytoplasmic contraction compared with wild-type cells (Fig. 3f). Given the role of MltG in termination^37^, we reasoned that the accumulation of synthases at the cell poles necessary for reducing cytoplasmic contraction was delayed in Δ*mltG* cells.

Taken together, these findings highlight the functional duality of the elongation machinery in both cell-wall synthesis and as a mechanically robust connection between the cytoplasmic membrane and the cell wall.

### The cytoplasmic membrane has substantial load-bearing capacity

Our findings collectively suggest that MreB does not directly contribute to envelope stiffness, and instead promotes physical coupling between the cytoplasmic membrane to the cell wall via its role in cell-wall synthesis. This linkage is unlikely to provide substantial stiffness in the longitudinal or circumferential directions along the surface of the cell, since the direction of forces exerted to link the layers would be predominantly perpendicular to the surface. However, if the cytoplasmic membrane itself were load bearing, such a coupling could explain the substantial decrease in bending rigidity under A22 treatment^20^, wherein MreB (and the coupling it promotes) is required for bending to engage the mechanical response of the cytoplasmic membrane (Fig. 4a). While it is typically assumed that the cytoplasmic membrane is relatively flexible compared to the cell wall by virtue of it being a lipid bilayer^12,38^, previous molecular dynamics simulations suggested that a protein-rich membrane patch mimicking the cytoplasmic membrane can exhibit substantial stiffness comparable to that of the cell wall and the outer membrane^39^.

**Figure 4:**
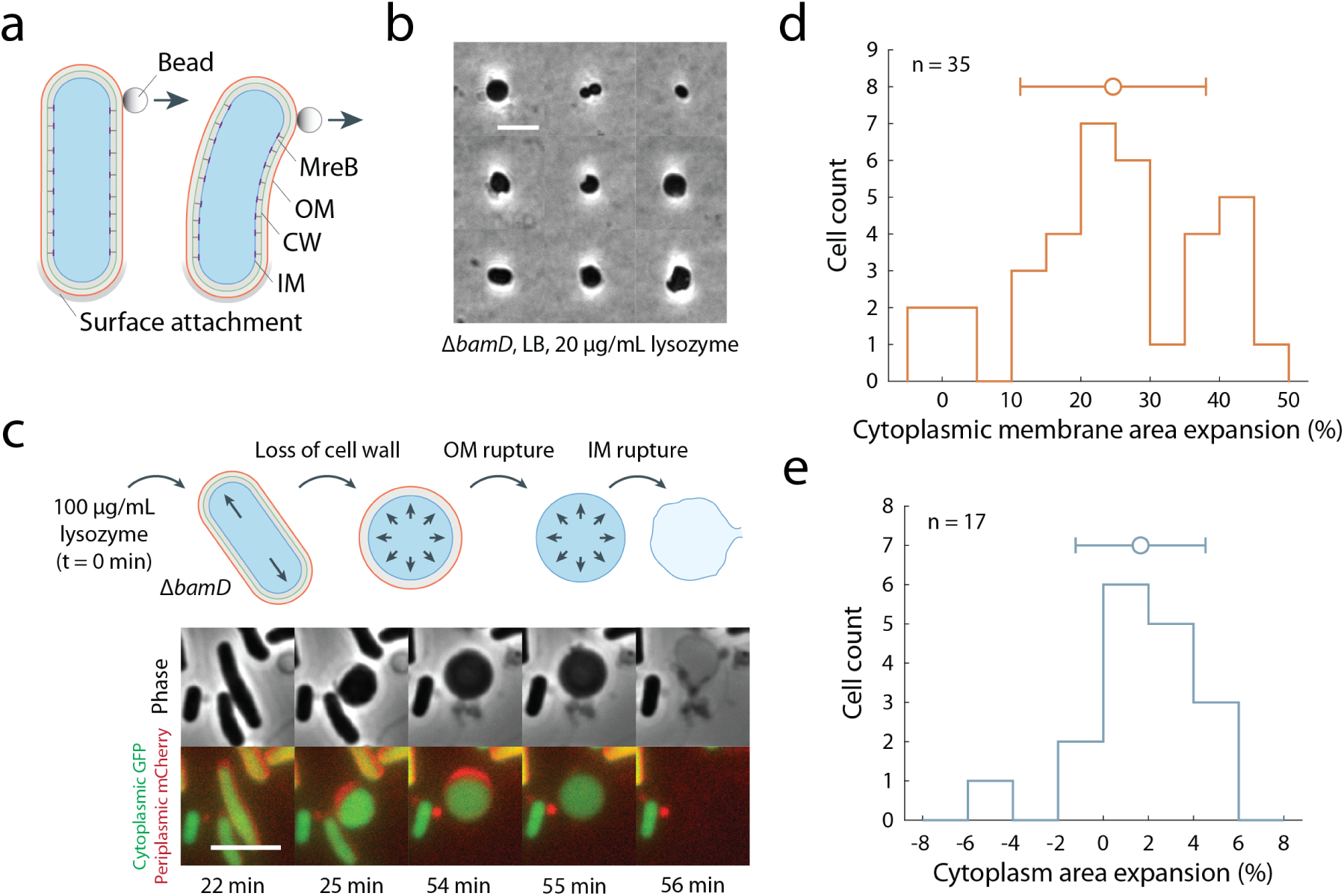
The cytoplasmic membrane has the capacity to bear substantial mechanical stress. a) MreB-mediated coupling between the cytoplasmic membrane and the cell wall may explain the decrease in bending rigidity of *E. coli* cells upon A22 treatment^20^ if the cytoplasmic membrane is sufficiently stiff. b) Spheroplasts without a cell wall can be generated from Δ*bamD* cells with weak outer membranes using 20 µg/mL lysozyme in LB medium without additional osmoprotection. Cells were imaged 3 h after lysozyme treatment (Methods). The panel shows a composite of representative snapshots of individual spheroplasts. Scale bar: 5 µm. c) Schematics and images during the generation of Δ*bamD* spheroplasts, rupture of the weakened outer membrane, and lysis due to inner membrane rupture. Δ*bamD* cells were labeled with cytoplasmic GFP and periplasmic mCherry and tracked in a microfluidic flow cell (Methods). Loss of periplasmic mCherry signal indicates rupture of the outer membrane, which always occurred prior to or at the same time as cytoplasmic membrane rupture. Scale bar: 5 µm. d) The surface area of the cytoplasmic membrane increased when the cell wall was degraded, indicating that the remaining envelope layers (cytoplasmic and outer membranes) bore more of the stress due to turgor pressure in the absence of the wall. *n*=35 cells. e) Rupture of the outer membrane in Δ*bamD* spheroplasts resulted in only a slight increase in cytoplasmic area, suggesting that the outer membrane was not bearing substantial stresses in the spheroplasts. *n*=17 cells.

To directly test the capacity of the inner membrane to bear stress, we sought to reduce or remove the stress-bearing capacity of the cell wall and outer membrane. Cell wall-free spheroplasts can be generated with lysozyme^40^. However, the outer membrane is thought to be essential, potentially due to osmotic balancing in the periplasm^17^, and no mechanism currently exists for robustly removing the outer membrane while retaining cell viability^12^. Thus, we leveraged our previous finding that deletion of *bamD* results in a highly mechanically compromised outer membrane due to a drastic reduction in outer membrane protein levels^41^. We successfully generated wall-free Δ*bamD* spheroplasts using 20 µg/mL lysozyme, indicating that even without a cell wall, at least some cells with a highly mechanically compromised OM can persist in a rich medium without osmoprotection (Fig. 4b).

To monitor changes to the cell envelope during and after cell-wall digestion, we tracked Δ*bamD* cells expressing cytoplasmic GFP and periplasmic mCherry in a microfluidic flow cell (Methods). Upon lysozyme introduction, cells lost their rod shape and became spherical (Fig. 4c). Without the stiff cell wall that typically bears turgor pressure, the mechanical load shifted to the two membranes, resulting in a ∼20% increase in the surface area of the cytoplasmic membrane (Fig. 4d, Methods). Approximately half of Δ*bamD* spheroplasts persisted for >10 min with an expanded cytoplasmic membrane before lysis, highlighting the ability of the membrane to bear turgor pressure. The spheroplasts survived for a shorter period compared to in a liquid environment (Fig. 4b), likely due to the mechanical confinement of the thin microfluidic flow cell. During the process of lysis, Δ*bamD* spheroplasts lost the periplasmic mCherry signal earlier than or simultaneously with the loss of cytoplasmic GFP (Fig. 4c), indicating that the outer membrane ruptured before or at the same time as the cytoplasmic membrane. After outer membrane rupture, ∼50% of spheroplasts persisted with an intact cytoplasmic membrane for >5 min. In these cells, outer membrane rupture caused almost no change in the cytoplasm area (Fig. 4e), suggesting that the turgor pressure was borne solely by the cytoplasmic membrane rather than the mechanically compromised outer membrane. These findings provide direct evidence of the load-bearing capability of the cytoplasmic membrane, such that when coupled to the cell wall by the process of cell-wall synthesis, the cytoplasmic membrane can increase the stiffness of the cell envelope.

## Discussion

In this study, we provide a revised perspective on the origins of mechanical resilience of the cell envelope, and demonstrate that the cell-wall synthesis machinery has a dual catalytic–structural function in integrating growth with envelope mechanics. We found that MreB does not contribute directly to cell wall stiffness under repeated plasmolysis (Fig. 1a,d) but instead functions as a platform for the components of the cell-wall synthesis machinery to link the cytoplasmic membrane and the cell wall sufficiently strongly as to prevent plasmolysis. Fluorescently labeled MreB filaments relocalized to the cell poles during large-magnitude osmotic shocks (Fig. 2), which in turn limited cytoplasmic contraction in these regions during periods of hyperosmolarity and led to peptidoglycan synthesis at the poles (Fig. 3b) as long as nutrients were available (Fig. 3a). Among the many components involved in cell-wall elongation, overexpression of RodA (Fig. 3c) and removal of MltG (Fig. 3e) impacted the connection between the cytoplasmic membrane and cell wall. Motivated by these findings and our evidence of substantial load-bearing capacity of the cytoplasmic membrane (Fig. 4), we propose a biophysical model in which MreB and active peptidoglycan synthesis are necessary for mechanically coupling the cytoplasmic membrane to the rest of the cell envelope. This model provides an explanation for the previously observed changes in bending rigidity during A22 treatment^20^: with intact MreB filaments, the cytoplasmic membrane can contribute to resisting bending. In fact, the ∼30% decrease in bending rigidity during A22 treatment is in line with expectations based on detaching a single layer from an envelope with three layers of roughly equal mechanical importance.

The proteins that physically link bacterial surface layers are often critical to cellular mechanical integrity. Braun’s lipoprotein (Lpp) and certain β-barrels connect the OM to the cell wall and impact the ability of the outer membrane to bear load under compression^13^. Lpp also regulates periplasmic width and the stiffness of *E. coli* cells^14^. However, the linkages between the cell wall and the cytoplasmic membrane have largely been overlooked, in part due to the lack of suitable assays for interrogating this connection. Our findings highlight the dynamic nature of the cytoplasmic membrane-peptidoglycan linkage. High-magnitude oscillatory osmotic shocks induced linkage formation at otherwise uncoupled cell poles, a process dependent on the relocalization of MreB and repositioning of at least certain components of the elongation machinery. The linkages, which are likely transient rather than persisting as static structures, may rely on several factors. While an obvious candidate is crosslinked glycans, our results suggest that, although they may contribute, they are not required. RodA does not extend far enough into the periplasm to bind peptidoglycan, suggesting that other factors such as RodZ or MreC mediate the connection. Potential redundancy or decoupling of the activity and binding of the factors involved may complicate elucidating the precise makeup and dynamics of the coupling. Nonetheless, it is striking that A22 treatment alone is sufficient to alter cytoplasmic contraction under repeated osmotic shocks (Fig. 1E), highlighting the key role of MreB in organizing the peptidoglycan synthesis machinery.

One open question raised by our work is the mechanism by which MreB relocalizes to the cell poles during large hyperosmotic shocks. Previous studies found that MreB is enriched in log-phase cells at regions of the membrane with negative mean curvature^33,42^, which has been hypothesized to underlie a feedback control mechanism for maintaining rod-like shape^9^. Plasmolysis generates regions of negative mean curvature near the poles, as indicated by the cytoplasmic membrane dye MitoTracker Green^43,44^ (Fig. S8a), and cytoplasmic contraction could generate wrinkles in the membrane with large absolute curvature that could either bias the localization of MreB filaments based on geometric preference or by disrupting their circumferential motion^45^. However, MreB relocalized to cell poles at a rate independent of whether obvious plasmolysis occurred at that pole (Fig. S8b), indicating that at least large deformations are not necessary for relocalization. Regardless of the mechanism, the ability of MreB to target distinct regions of the cell depending on environmental cues such as osmolarity may provide bacterial cells with morphological flexibility. This flexibility is apparent during the transition into and out of stationary phase, during which MreB also exhibits polar localization^24^, which our study suggests could be due to osmolarity changes.

In addition to mechanically strengthening the cell envelope, linkages between surface layers control the distance between layers^14^, which can be critical for the function of proteins that span the space^46–48^. In the case of MreB, the cell-wall synthesis-mediated linkages are mostly present only along the cylindrical cell body during log-phase growth. Thus, during a small osmotic perturbation, the cell poles can function as a temporary buffer for water efflux, while the distance between the cytoplasmic membrane and the cell wall along the rest of the cylindrical body is not affected, allowing for continued growth even in the absence of turgor^13,49^. In the future, it will be interesting to determine whether MreB-directed cell-wall synthesis plays a similar mechanical function in other species. Regardless, a deeper understanding of the mechanical interactions between the cytoplasmic membrane and the cell wall provides new insights into the structural stability of the cell envelope and reveals potential targets and mechanisms for antimicrobial intervention.

## Methods

### Bacterial strains and growth conditions

All strains and plasmids used in this study are listed in Table S1. *E. coli* cells were grown overnight in lysogeny broth (LB, Lennox, Fisher Scientific, Cat. #BP9722-500) at 37 °C, diluted into fresh LB, and grown for 2 h into log phase for microfluidic measurements. Strains were grown with 50 µg/mL kanamycin (Sigma-Aldrich, Cat. #K4000) or 50 µg/mL chloramphenicol (Calbiochem, Cat. #220551) as necessary.

The Δ*mltG* strain expressing cytoplasmic GFP was generated by λ-Red recombination using pSIM6. A chloramphenicol resistance cassette (cmR) flanked by FRT sites was PCR-amplified from pKD3 with primers containing the same 50-nt homology overhangs upstream and downstream of *mltG* as the Keio collection^50^ (forward primer: 5’-ACGTTATATGAATATTTAGCCCCACTTTGTGAGCGCCCGAATTAGTCATGtccatatgaatatcctccttagttcctattcc-3’; reverse primer: 5’-CGCCTTCCAGCCCCTCAATGACGATATACTTACTGCGCATTTTTTTCCTTtgtaggctggagctgcttcg-3’). The PCR product was electroporated into BW25113 carrying pSIM6, and recombinants were selected on LB agar with 50 µg/mL chloramphenicol. The strain was subsequently transformed with pZS21-GFP (Table S1) to drive constitutive GFP expression.

### Microfluidic hyperosmotic shock assays

Cells from a saturated culture grown overnight were diluted 1:200 into fresh LB and grown at 37 °C for 2 h. These log-phase cells were cultured with 25 µg/mL wheat germ agglutinin (WGA) conjugated to AlexaFluor-555 (AF555, Invitrogen W32464) to label the outer membrane. Cells were then loaded into CellASIC ONIX microfluidic flow cells (Sigma-Aldrich, Cat. #B04A-03-5PK), in which media were exchanged using the CellASIC ONIX2 microfluidic platform (Sigma-Aldrich, Cat. #CAX2-S0000). Oscillatory hyperosmotic shock assays were performed at room temperature.

Cells were first perfused with LB for 30 min with 50 µg/mL cephalexin (MP Biomedicals, Cat. #150585) to inhibit cell division and then treated with 100 µg/mL A22 (gift from Douglas Weibel), 80 µg/mL MP265 (MedChemExpress, Cat. #HY-131583), a beta-lactam antibiotic (100 µg/mL ampicillin (RPI, Cat. #A40040), 50 µg/mL mecillinam (Sigma-Aldrich, Cat. #33447), or 30 µg/mL cefsulodin (MP Biomedicals, Cat. #198677), or 0.85X PBS (Gibco, Cat. #70011044) for 10 min to perturb cell-wall synthesis. Cells were then subjected to oscillatory hyperosmotic shocks involving 400-, 600-, or 1000-mM sorbitol (Sigma-Aldrich, Cat. #S1876, dissolved in LB or 0.85X PBS) and recovery without sorbitol, with a period of 4 min (2 min shock and 2 min recovery). Pre-treatment compounds (A22, MP265, beta-lactam, or 0.85x PBS) were maintained during osmotic shocks. Cells were stained with 25 µg/mL WGA-AF555 throughout incubation in the flow cell.

To overexpress RodA or RodA^D262N^ (Table S1), cells were induced with 1 mM IPTG (Sigma-Aldrich, Cat. #I6758) for 1 h in liquid culture before loading to the microfluidic flow cell. After loading, cells were incubated with 1 mM IPTG for an additional 30 min, and IPTG was maintained throughout subsequent osmotic shock assays.

### Fluorescent labeling of peptidoglycan and cytoplasmic membrane

A saturated culture grown overnight in LB medium was diluted 1:200 into LB medium containing 1 mM HADA (MedChemExpress, Cat. #HY-131045/CS-0124027) and grown at 37 °C for 2 h to ensure homogenous peptidoglycan labeling. Cells with fluorescently labeled peptidoglycan were then loaded into microfluidic flow cells and grown in LB with 1 mM HADA at room temperature, followed by hyperosmotic shocks with 1 mM HADA. Cells were then switched to HADA-free LB medium to track peptidoglycan synthesis as indicated by the decay of HADA signals.

To visualize the cytoplasmic membrane during osmotic shocks, cells were labeled with 2 µg/mL MitoTracker Green (Invitrogen, Cat. #M7514) in liquid culture and in the microfluidic flow cell.

### Single-cell imaging

Cells undergoing oscillatory hyperosmotic shocks were monitored using a Nikon Eclipse Ti-E inverted epifluorescence microscope with a 100X (NA 1.40) oil-immersion objective (Nikon Instruments). Phase-contrast and fluorescence images were collected on a Prime BSI Express sCMOS camera (Teledyne Photometrics) using *µManager* v. 2.0^51^.

### Image analysis

Time-lapse images were segmented with the machine learning-based software *DeepCell*^52^ trained on WGA-stained cells, and the segmented images were analyzed using *Morphometrics*^52^ to obtain cell contours at sub-pixel resolution. Using these contours, single-cell lineages were tracked. Cells that were segmented or tracked incorrectly or that lysed during osmotic shocks were filtered out via manual inspection. Cell length and width were calculated using the *MicrobeTracker*^53^ meshing algorithm. Cytoplasmic area was inferred using cytoplasmic GFP fluorescence, and cytoplasmic contraction during plasmolysis was computed based on the fraction of the total area enclosed by the cell contour represented by the cytoplasmic area.

To determine the spatial localization of MreB filaments in cells expressing MreB-GFP, in Fig. 2b, fluorescence profiles along cell contours were calculated as previously described^54^ and the degree of fluorescence along the cylindrical region and the cell poles was compared. In Fig. 2c, S4, and S6b, the average fluorescence intensity was calculated for the projected cylindrical region and both cell poles, where the highly curved polar regions were defined based on a curvature threshold.

### Cell shape analysis

Cell wall area contraction in Fig. 1d was calculated as (area plasmolysis contraction)/(area after plasmolysis)–1. Cytoplasmic area expansion in Fig. 4e was calculated as (area after OM rupture)/(area before OM rupture)–1. Cell width was extracted from the pill mesh generated by the *MicrobeTracker*^53^ meshing algorithm and defined as the maximum width measured perpendicular to the cell’s long axis. Cell width increase in Fig. S3b was calculated as (width under shock)/(width prior to shock)–1.

The cytoplasmic membrane area in Fig. 4d was calculated separately for rod-shaped cells prior to wall digestion and for spheroplasts after digestion using the same approximation as in ref. ^13^. Cell contours were generated using *Morphometrics*^52^ from cytoplasmic GFP fluorescence. For rod-shaped cells, the membrane area was estimated from the contour assuming rotational symmetry around the long axis. For spheroplasts, which adopted amorphous shapes within the thin flow cell, the flow cell height was assumed equal to the cell width prior to wall digestion, *w*. The projected area, *a*, and perimeter, *s*, were calculated from the contour, and the cytoplasmic membrane area was then estimated as 2*a*+*ws*.

### Spheroplast generation

To generate cell wall-free spheroplasts with a compromised OM, Δ*bamD* cells (Table S1) were grown in LB for 2 h and treated with 100 µg/mL lysozyme (Sigma-Aldrich, Cat. #L6876) in LB. To generate and monitor spheroplasts in a microfluidic flow cell, Δ*bamD* cells were treated with 100 µg/mL lysozyme in LB. Expression of periplasmic mCherry was induced with 1 mM IPTG.

## Data and code availability

All data used for generating figures in this study are available at the Stanford Digital Repository: https://purl.stanford.edu/gg063dh4581.

## Supplementary Figures

**Figure S1:**
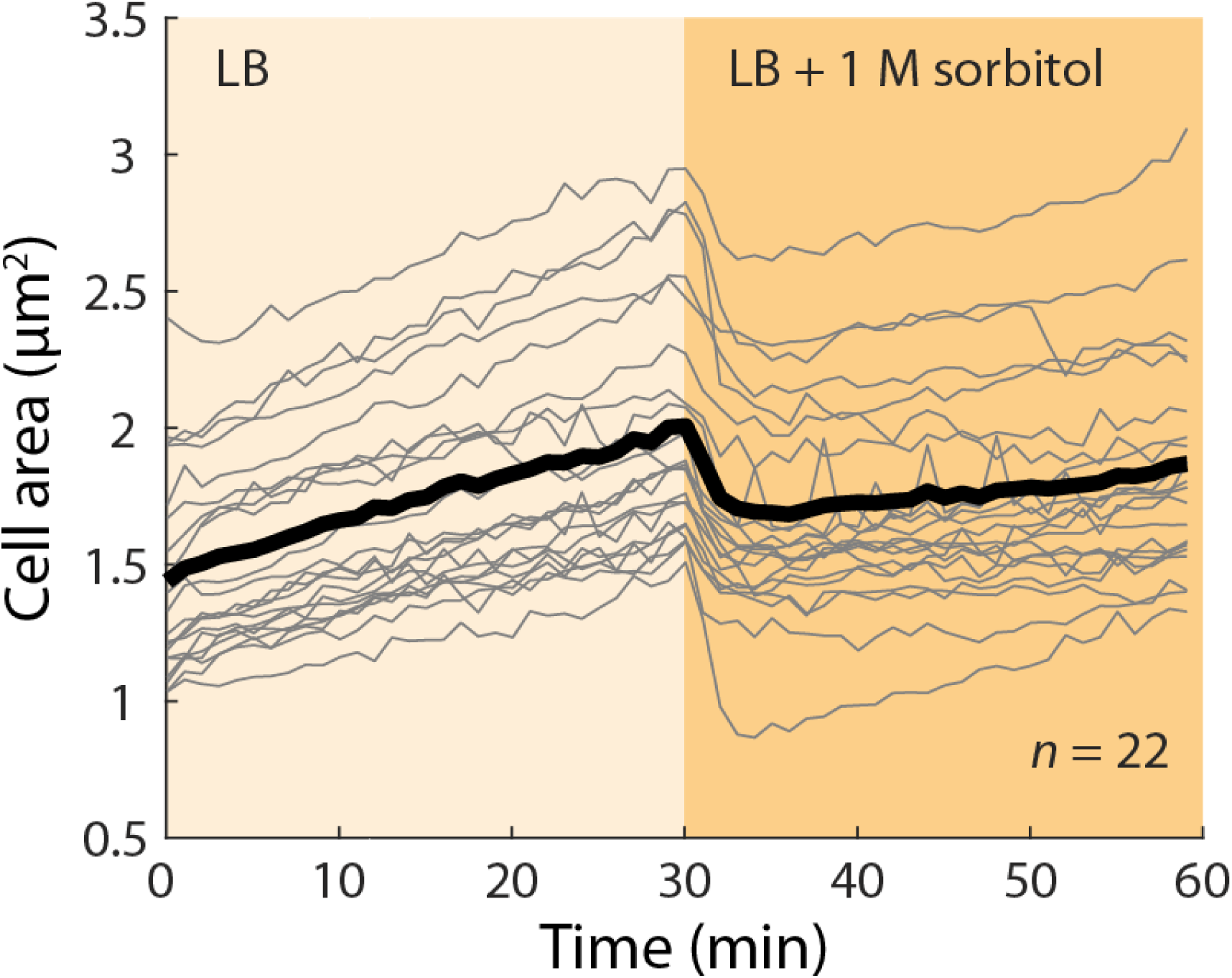
Cell growth is inhibited under large hyperosmotic shocks. During a hyperosmotic shock with 1 M sorbitol in LB, cell growth was inhibited for an extended period. Cells were monitored in a microfluidic flow cell (Methods). *n*=22 cells.

**Figure S2:**
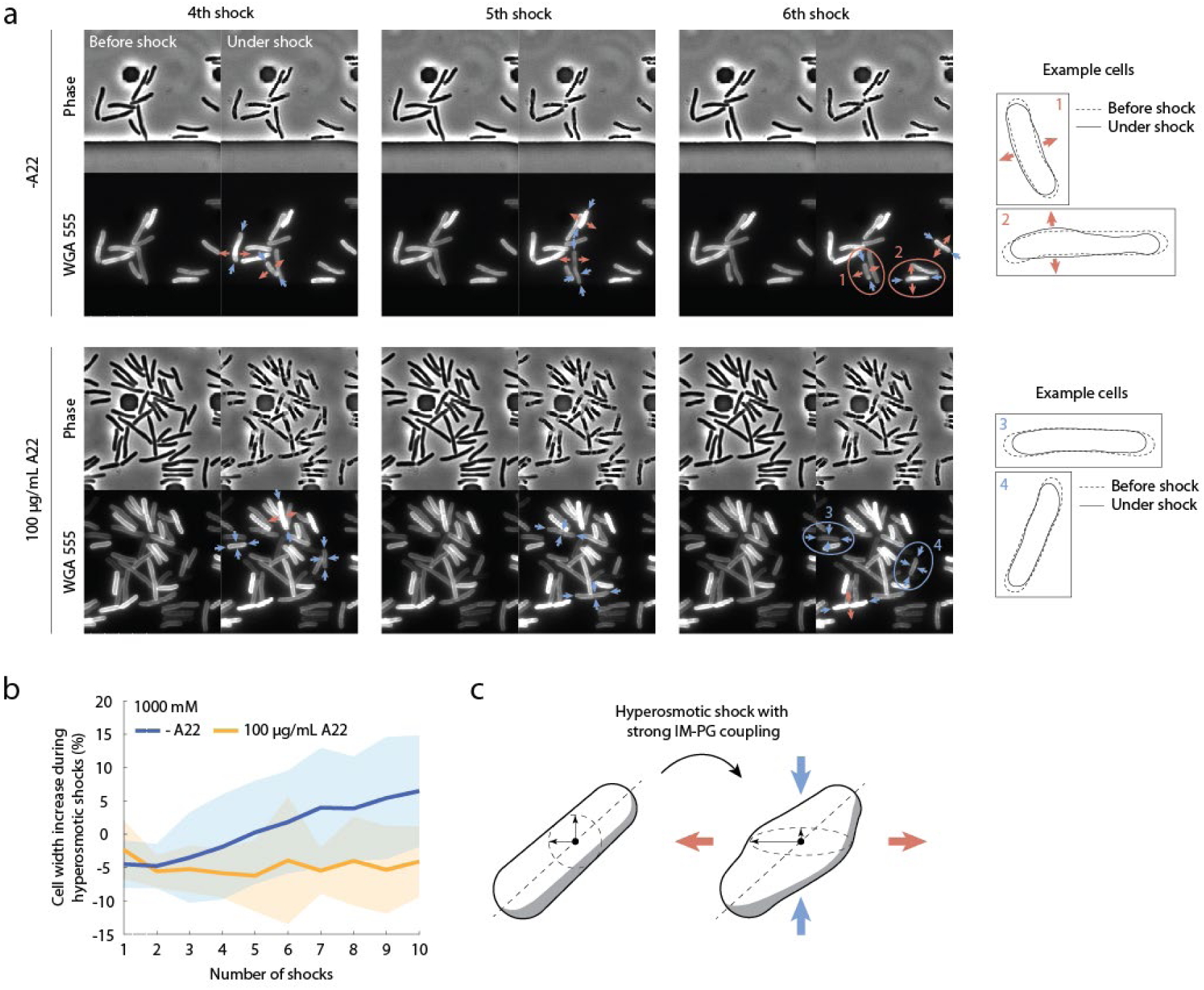
When mechanically coupled to the cytoplasm, the cell wall exhibits large deformations and increases in cell width during large hyperosmotic shocks. a) After several osmotic shock cycles (1 M sorbitol in LB), when the cell wall is mechanically coupled with the cytoplasmic membrane by MreB filaments, the cell wall becomes flattened by the negative osmotic pressure from inside the cytoplasm under high environmental osmolarity, resulting in increased cell width around midcell. These large deformations only occurred under large (600 mM or 1 M) osmotic shocks. b) Cell width increased after several 1-M hyperosmotic shock cycles. This widening was abolished by A22 pre-treatment, with cells contracting in width upon loss of turgor. c) Schematic of cell flattening caused by the mechanical linkages and osmotic pressure. Red and blow arrows indicate extension and compression along different axes of the cross-section.

**Figure S3:**
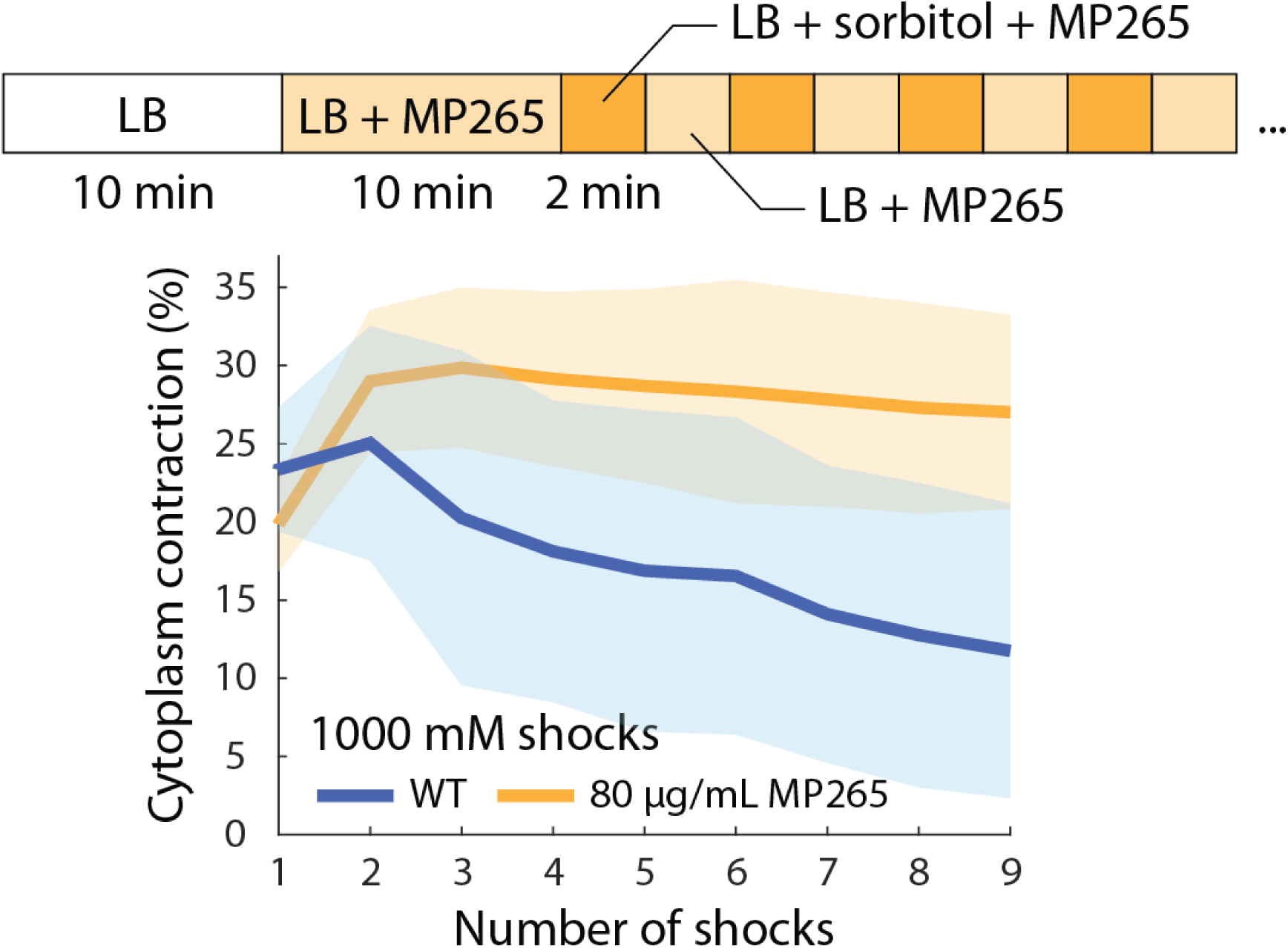
Disruption of MreB filaments with MP265 eliminates the mechanical coupling between the cytoplasmic membrane and the cell wall during oscillatory osmotic shocks. Wild-type cells were treated with 80 µg/mL MP265 10 min prior to and during 1-M osmotic shocks. The lack of coupling is similar to the effects of A22 treatment (Fig. 3e).

**Figure S4:**
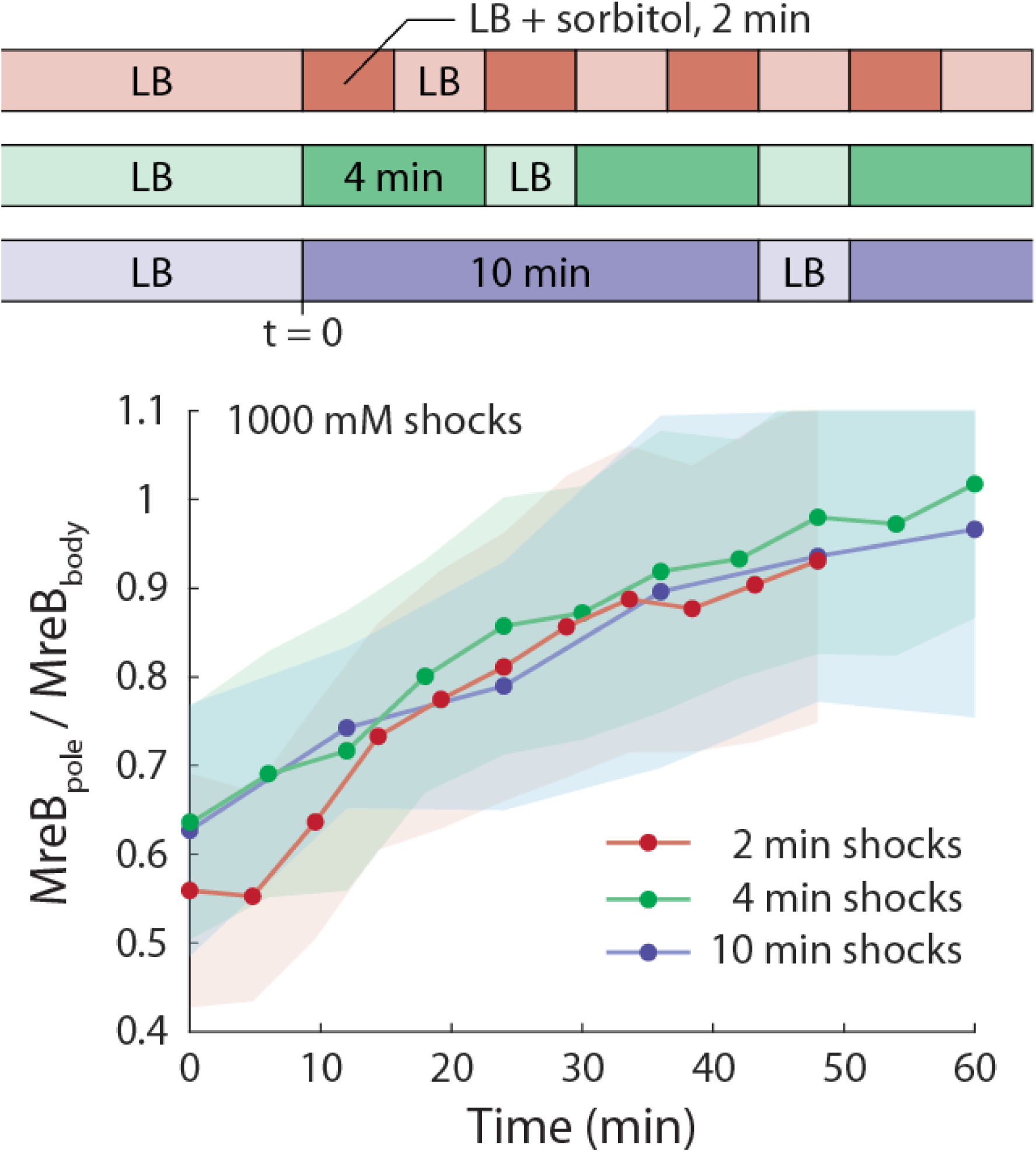
MreB relocalization to the poles is determined by the time under osmolarity oscillation rather than the number of shock cycles. Hyperosmotic shocks with extended periods (4 min, 10 min) of hyperosmolarity (upper schematic) resulted in similar degrees of MreB relocalization as with a 2-min period for a given assay time. Data points represent individual osmotic shock cycles. Fluorescence was evaluated during times of normal osmolarity to avoid bias caused by plasmolysis.

**Figure S5:**
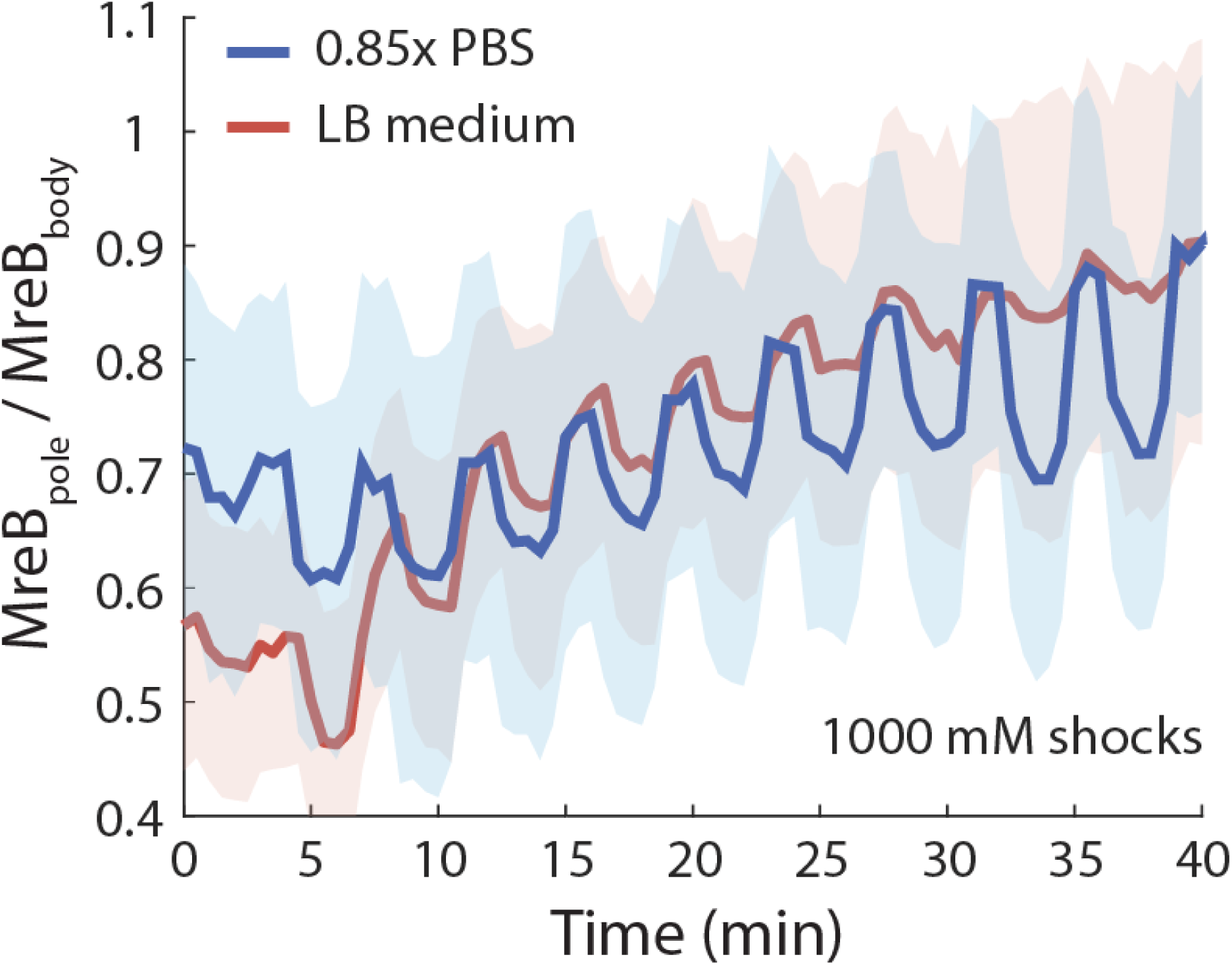
MreB relocalizes to the poles during oscillatory osmotic shocks in a nutrient-free buffer. In 0.85X PBS, MreB relocalization due to osmotic shocks progressed at a similar rate as in LB.

**Figure S6:**
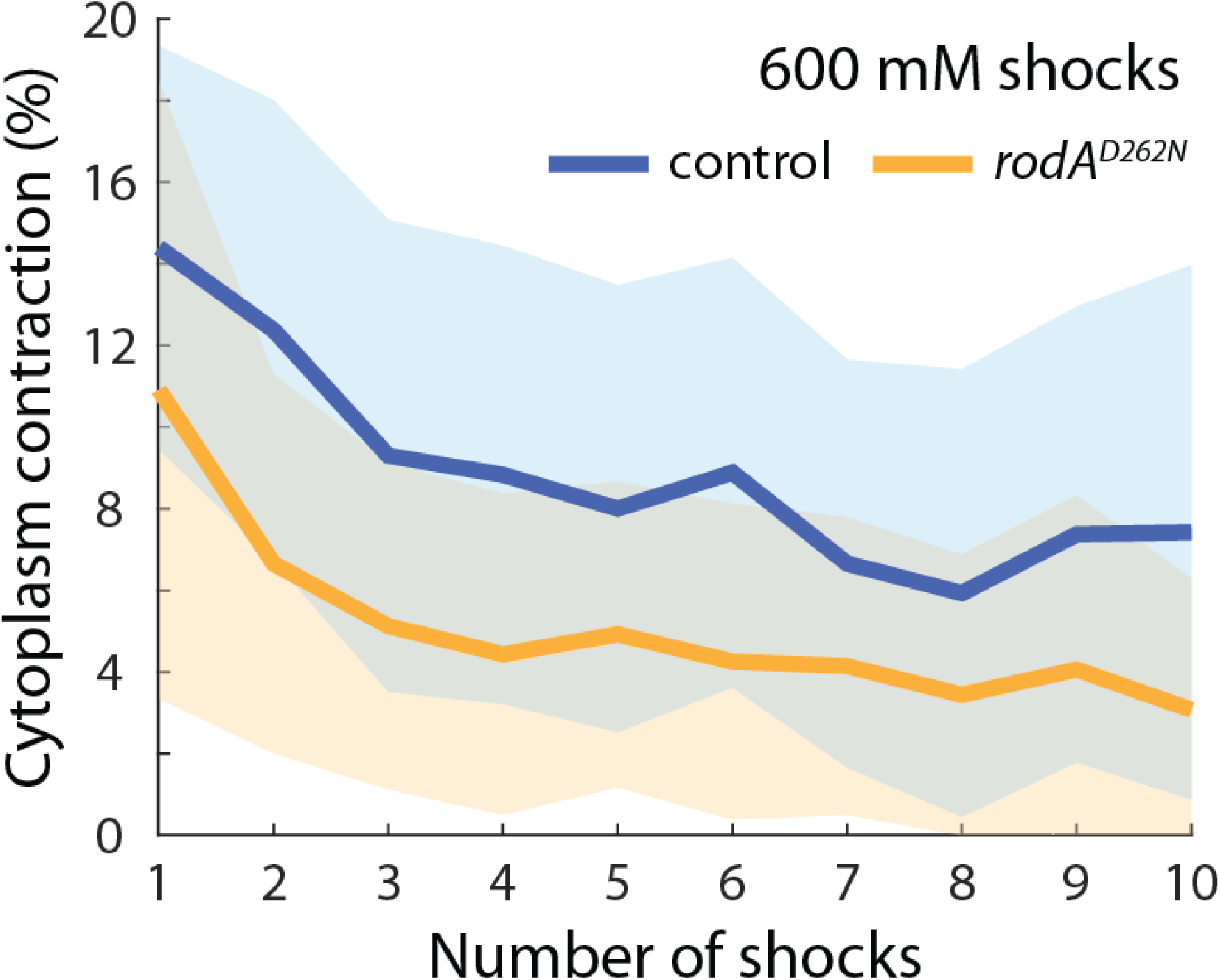
Overexpression of the dominant negative mutant *rodA^D262N^* reduced cytoplasmic contraction during 600-mM osmotic shocks.

**Figure S7:**
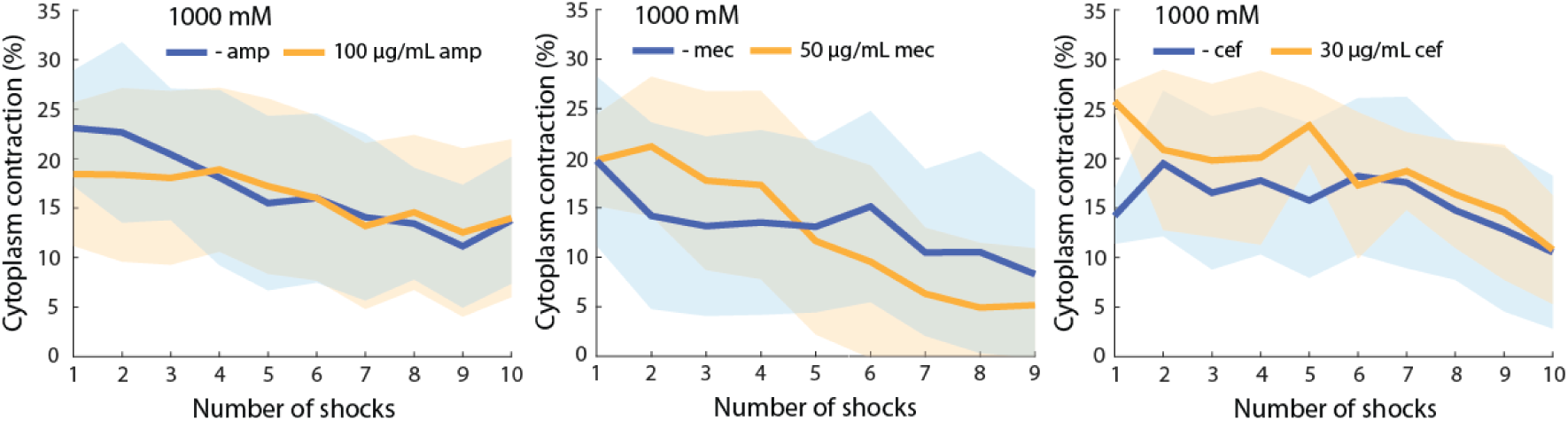
Inhibition of PBPs by 50 µg/mL mecillinam, 30 µg/mL cefsulodin, or 100 µg/mL ampicillin did not impact the dynamics of cytoplasmic contraction during 1-M shocks.

**Figure S8:**
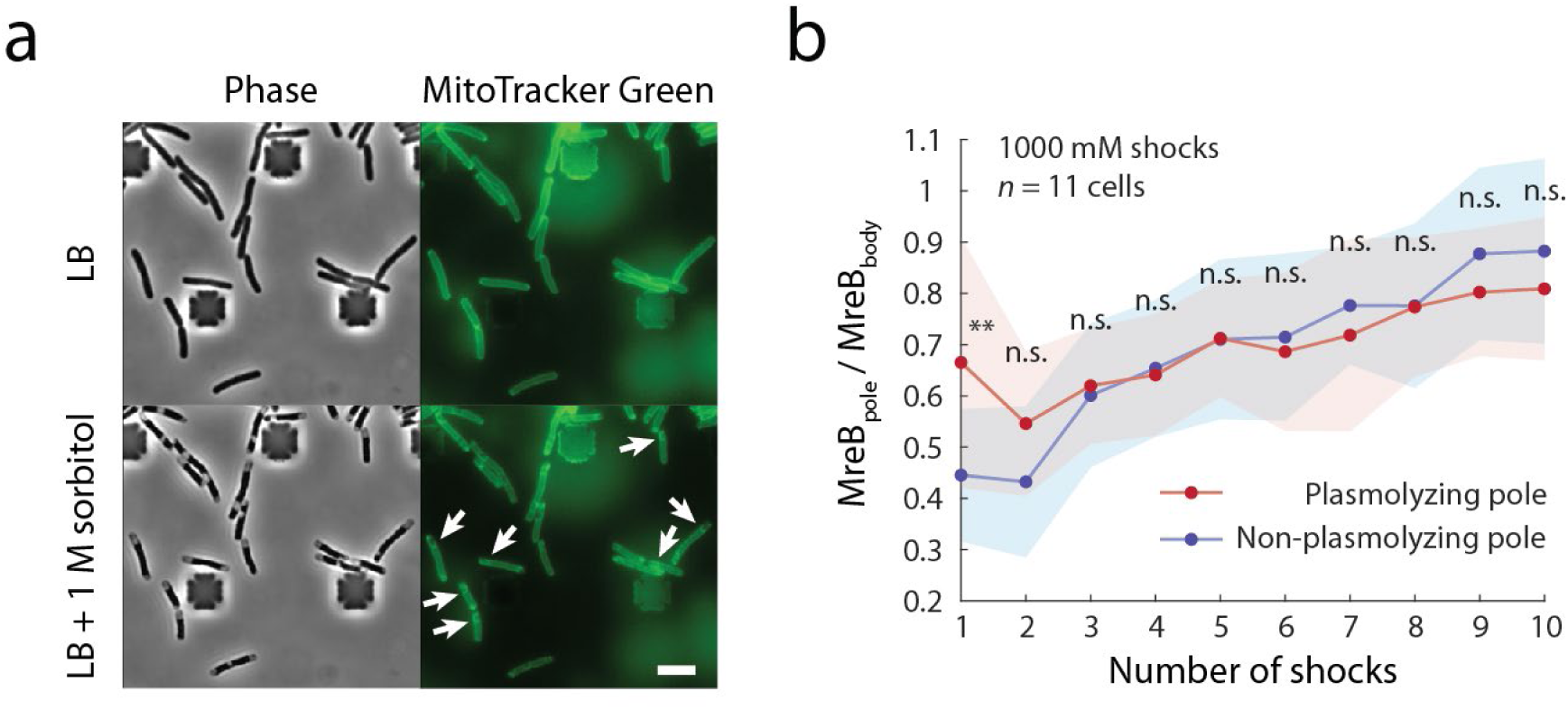
MreB relocalization to each cell poles during hyperosmotic shocks is not dependent on the magnitude of cytoplasmic contraction at the pole. a) Plasmolysis resulted in clear regions with negative mean curvature near cell poles. The cytoplasmic membrane was stained with MitoTracker Green. Scale bar: 5 µm. b) During plasmolysis, many cells exhibited cytoplasmic contraction at only one cell pole. No significant difference was observed between MreB relocalization at the two poles after repeated shocks. Fluorescence was evaluated during times of normal osmolarity to avoid bias caused by plasmolysis. *n*=11 cells. **: *p* < 0.01, n.s.: *p* > 0.01; Welch’s t-test was performed for each osmotic shock cycle.

## Supplementary Table

**Table S1:** *E. coli* strains and plasmids used in this study.

| Strain | Genotype | Source/reference |
| --- | --- | --- |
| MG1655 | F<sup>-</sup>, lambda<sup>-</sup>, *rph-1* | CGSC #6300 |
| KC1016 | MG1655 pZS21-GFP | Gift from Thomas Silhavy |
| KC508 | MG1655 *mreB-msfGFP* | 33 |
| KC496 | MG1655 $\Delta lacIZYA::frt \Delta ponA::frt$<br>$att\lambda::pTB309(Para::ponA)$ | 55 |
| pPR1 | MG1655 *Ptac::pbpA-rodA*<sup>D262N</sup> | 35 |
| pPR6 | MG1655 *Ptac::pbpA-rodA* | 35 |
| BW25113 | F<sup>-</sup> $\lambda^- \Delta(araD-araB)567$<br>$\Delta lacZ4787(::rrnB-3) rph-1 \Delta(rhaD-$<br>$rhaB)568 hsdR514$ pZS21-GFP | Gift from Jie Xiao |
| $\Delta mltG$ | BW25113 $\Delta mltG::cmR$ pZS21-GFP | This study |
| IMB1056 | MC4100 *ara<sup>R</sup> bamA*<sup>E470K</sup> $\Delta bamD$ | 41 |
| IMB1144 | IMB1056 *attHK::Plac-mCherry*<br>pZS21-GFP | 41 |
| Plasmids | Genotype | Source/reference |
| pZS21-GFP | *pZS21kan::GFP* | Gift from Thomas Silhavy |
| pPR1 | *Ptac::pbpA-rodA*<sup>D262N</sup> | 35 |
| pPR6 | *Ptac::pbpA-rodA* | 35 |

## Supplemental Movies

**Supplemental Movie 1: A22 treatment (100 µg/mL) depolymerizes MreB filaments within 10 min in LB medium.** Scale bar: 5 µm.

## Acknowledgements

The authors thank the Huang lab for helpful discussions. The authors acknowledge support from a Stanford Interdisciplinary Graduate Fellowship (to J.S.), NIH RM1 Award GM135102 (to K.C.H.), and NSF Award EF-2125383 (to K.C.H.). K.C.H. is a Chan Zuckerberg Biohub Investigator. This work was also supported in part by the National Science Foundation under Grant PHYS-1607611 and the hospitality of the Aspen Center for Physics.

## Notes

https://purl.stanford.edu/gg063dh4581

### Competing Interest Statement

The authors have declared no competing interest.

### Summary of Updates

The figures have been updated to higher-resolution versions.

